# Structural diversity and oligomerization of bacterial ubiquitin-like proteins

**DOI:** 10.1101/2024.11.21.623966

**Authors:** Minheng Gong, Qiaozhen Ye, Yajie Gu, Lydia R. Chambers, Andrey A. Bobkov, Neal K. Arakawa, Mariusz Matyszewski, Kevin D. Corbett

**Affiliations:** Department of Cellular and Molecular Medicine, University of California San Diego, La Jolla CA 92093; Department of Chemistry and Biochemistry, University of California San Diego, La Jolla CA 92093; Conrad Prebys Center for Chemical Genomics, Sanford Burnham Prebys Medical Discovery Institute, CA 92037; Environmental and Complex Analysis Laboratory, Department of Chemistry and Biochemistry, University of California San Diego, La Jolla CA 92093; UC San Diego Cryo-EM Facility, University of California San Diego, La Jolla CA 92093; Department of Molecular Biology, University of California San Diego, La Jolla CA 92093

## Abstract

Bacteria possess a variety of operons with homology to eukaryotic ubiquitination pathways that encode predicted E1, E2, E3, deubiquitinase, and ubiquitin-like proteins. Some of these pathways have recently been shown to function in anti-bacteriophage immunity, but the biological functions of others remain unknown. Here, we show that ubiquitin-like proteins in two bacterial operon families show surprising architectural diversity, possessing one to three β-grasp domains preceded by diverse N-terminal domains. We find that a large group of bacterial ubiquitin-like proteins possess three β-grasp domains and form homodimers and helical filaments mediated by conserved Ca^2+^ ion binding sites. Our findings highlight a distinctive mode of self-assembly for ubiquitin-like proteins, and suggest that Ca^2+^-mediated ubiquitin-like protein filament assembly and/or disassembly enables cells to sense and respond to stress conditions that alter intracellular metal ion concentration.

## Introduction

Ubiquitination is a multi-step enzymatic cascade in which ubiquitin or a ubiquitin-like protein (Ubl) becomes covalently linked to a target - typically a lysine side chain or other amine group - through the sequential action of E1, E2, and (usually) E3 proteins^1^. Ubiquitination and related pathways regulate protein homeostasis and other processes in eukaryotes^2–6^, but related bacterial pathways are usually involved in metabolic pathways and do not mediate protein conjugation. Recently, bioinformatics analysis of bacterial genomes has identified several sparsely distributed families of bacterial operons that encode proteins related to eukaryotic E1, E2, E3, Ubl, and peptidase/deubiquitinase proteins^7–9^. Structural and biochemical studies of three such operon families have now demonstrated that they are related to eukaryotic ubiquitination pathways and perform *bona fide* protein conjugation reactions^10–13^. Type II CBASS anti-bacteriophage (phage) operons encode an E2-E1 fusion protein, Cap2, that conjugates the C-terminus of their cognate CD-NTase to unknown targets to activate CBASS signaling^10,11^. Type I and II Bil (Bacterial ISG15-like) operons encode separate E1, E2, Ubl, and DUB proteins, and when cells are infected with phages, these pathways conjugate their Ubl to a phage tail protein to inhibit virion assembly and infectivity^12,13^. Finally, in an accompanying work, we find that <u>B</u>acterial <u>ub</u>iquitination-like (Bub) operons, previously termed “6E” or “DUF6527” operons, also encode E1, E2, Ubl, and peptidase proteins and perform protein conjugation^14^. These studies show that ubiquitination pathways evolved and proliferated in bacteria in the contexts of antiphage immunity and stress response.

In eukaryotes, Ubls typically comprise a single small domain with a fold termed “β-grasp”^1^. Some Ubls, notably including the innate-immune protein ISG15, possess two β-grasp domains. In prior work, we found that bacterial Ubls in Type II Bil operons show high structural diversity, with up to three predicted β-grasp domains and diverse fused N-terminal domains^12^. Here, we expand on that finding and show that Ubls in Bil and Bub operons possess diverse architectures, and that many such proteins form higher-order oligomers. We find that a large subset of bacterial Ubls contain three β-grasp domains and form filamentous assemblies *in vitro* upon calcium (Ca^2+^) ion binding. We show that Ca^2+^-induced filament formation occurs in diverse Ubls from Type II Bil, Type I Bub, and Type II Bub operons, suggesting that this property plays an important role in Bil/Bub operon function. We propose a mechanism in which assembly and/or disassembly of Ubl filaments enables cells to respond to changes in metal ion concentration during phage infection and/or other stress conditions.

## Results

### Bacterial ubiquitin-like proteins are architecturally diverse

To comprehensively characterize the architectural diversity of bacterial Ubls, we took advantage of recent studies that identified hundreds of bacterial operons encoding different combinations of ubiquitination-related proteins, particularly those that encode a predicted Ubl. These operons fall into two major groups termed Bil (<u>B</u>acterial ISG15-like)^15^ and Bub (<u>B</u>acterial <u>ub</u>iquitination-like) operons^14^. Bil operons can be divided into two families, Type I and Type II, based on the sequences of their proteins. All Bil operons encode a Ubl (BilA), an E2 (BilB), a JAB/JAMM-family peptidase/DUB (BilC), and an E1 protein (**Figure 1a**)^12,15^. Bub operons can also be divided into two families, which encode different but related domains. Type I Bub operons encode a Ubl-E2 fusion (BubAB), a JAB/JAMM peptidase-E1 fusion (BubCD), and a DUF6527 protein with serine protease activity (BubE; **Figure 1a**)^14^. Type II Bub operons encode a Ubl (BubA), an E2-E1 fusion (BubBD), and a DUF6527 protein (BubE), and typically do not encode a JAB/JAMM peptidase (**Figure 1a**).

**Figure 1.**
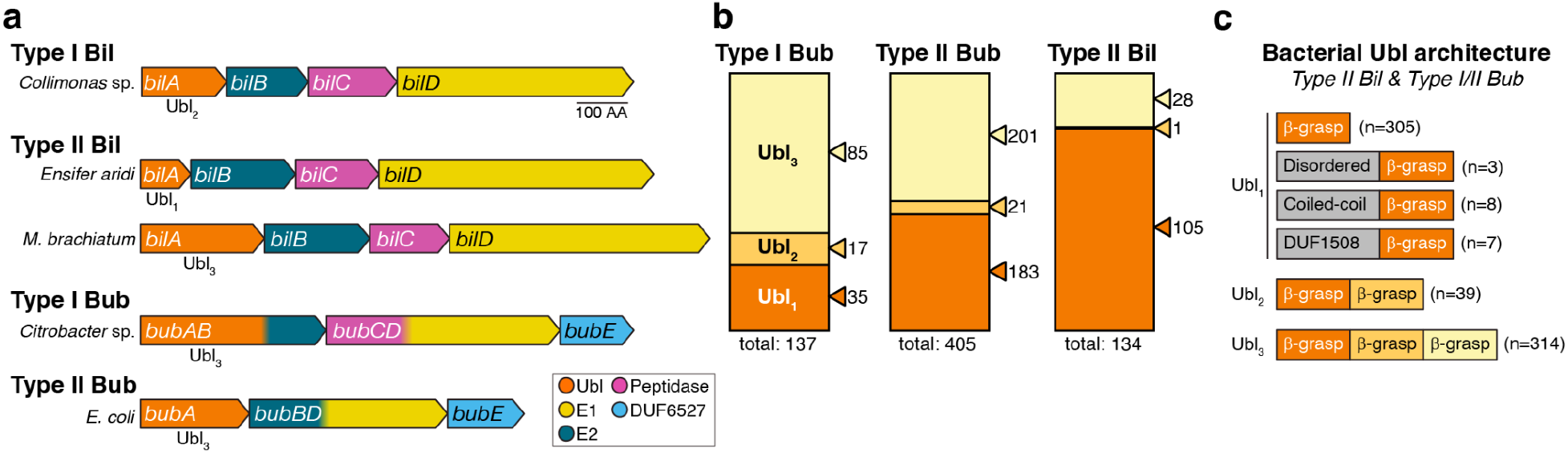
Bacterial ubiquitin-like proteins show diverse architectures. **(a)** Operon schematics of representative Type I and Type II Bil (Bacterial ISG15-like) operons, and Type I and Type II Bub (Bacterial ubiquitination-like) operons. Ubiquitin-like genes (*bilA/bubA*) are shown in orange; E2-like genes (*bilB/bubB*) in dark blue, JAB/JAMM peptidase genes (*bilC/bubC*) in magenta, E1-like genes (*bilD/bubD*) in yellow, and DUF6527 genes (*bubE*) in light blue. See **Table S1** for a catalog of Type II Bil, Type I Bub, and Type II Bub operons. **(b)** Ubl architecture in Type I Bub, Type II Bub, and Type II Bil operons. Ubls with one β-grasp domain (Ubl_1_) are shown in dark orange, those with two β-grasp domains (Ubl_2_) light orange, and three β-grasp domains (Ubl_3_) in light yellow. See **Table S1. (c)** Summary of Ubl architecture in Type I Bub, Type II Bub, and Type II Bil operons. Ubl_1_ proteins show diverse N-terminal domains. See **Figure S1** for structural predictions of Ubl_1_ proteins with predicted disordered, coiled-coil, or DUF1508 N-terminal domains.

We used protein sequence alignments and structure predictions from ESMFold^16^ and AlphaFold^17,18^ to comprehensively outline the predicted domain architectures of Ubls in previously-identified Bil and Bub operons (**Figure 1b-c, Table S1**)^15^. Type I Bil operons overwhelmingly encode Ubls with two predicted β-grasp domains, similar to the structure of ISG15 (ref. ^15^), and are not further considered here. Ubls in Type II Bil operons show diverse predicted architectures: 105 out of 134 identified operons (78%) encode a Ubl with a single predicted β-grasp domain, 1 (<1%) contains two β-grasp domains, and 28 (21%) contain three β-grasp domains (**Figure 1b**). Some Ubls with a single predicted β-grasp domain possess a variable N-terminal domain; we identified one Ubl from *Hymenobacter coccineus* with a 151 amino acid N-terminal tail rich in asparagine and glycine (N/G), suggesting that this protein may undergo liquid-liquid phase separation (**Figure S1a-c, Table S1**). We also identified five Ubls with a predicted N-terminal coiled-coil domain, suggesting that these proteins may form homodimers (**Figure S1f-i, Table S1**).

As in Type II Bil, Ubls from Type I and II Bub operons Ubls show diverse predicted architectures: 218 (40%) contain a single predicted β-grasp domain, 38 (7%) contain two β-grasp domains, and 286 (53%) contain three β-grasp domains (**Figure 1b**). Among those Ubls with a single predicted β-grasp domain, we identified two with long N/G-rich N-terminal tails, and three with predicted N-terminal coiled-coils (**Table S1**). We also identified 7 Ubls with tandem DUF1508 domains followed by a single β-grasp domain (**Figure 1c, Figure S1d-e**). DUF1508 domains are found in stress-induced genes including *E. coli* YegP^19,20^ and appear either as single domains that form homodimers (PDB ID 6Q2Z)^20^, tandem domains that form symmetric pseudo-dimers (PDB 2K8E), or as extended arrays (e.g. Pfam PF07411). Overall, our sequence analyses show that bacterial Ubls from Type II Bil and Type I/Type II Bub operons possess diverse architectures with variable numbers of β-grasp domains, plus other domains that could mediate self-association or oligomerization (**Figure 1c**). Because all of these proteins are named either “BilA” or “BubA” but these names do not capture their structural diversity, in the following sections we use a standard nomenclature with “Ubl” followed by a subscript number indicating the number of β-grasp domains in each protein (e.g. “Ubl_3_” for a protein with three predicted β-grasp domains).

We performed size exclusion chromatography coupled to multi-angle light scattering (SEC-MALS) on six Ubl proteins with different predicted architectures: two encoding two predicted β-grasp domains (Ubl_2_), two with a predicted coiled-coil domain followed by one β-grasp domain (CC-Ubl_1_), and two with N-terminal tandem DUF1508 domains followed by one β-grasp domain (DUF1508-Ubl_1_). The Ubl_2_ proteins and the DUF1508-Ubl_1_ proteins all formed monomers in solution (**Figure S2a-d**). We observed that one CC-Ubl_1_ formed a homodimer, while the other was monomeric (**Figure S2e-f**).

### Structures of Ubl_3_ proteins reveal a conserved homodimeric assembly

Bacterial Ubls predicted to contain three β-grasp domains (Ubl_3_) are nearly as common as those encoding one β-grasp domain. We expressed and purified three diverse Ubl_3_ proteins from a Type II Bil operon (*Methylobacterium brachiatum* DSM 19569 Ubl_3_), a Type I Bub operon (*Citrobacter* sp. RHBSTW-00271 Ubl_3_), and a Type II Bub operon (*E. coli* ZDHYS365 Ubl_3_). We first crystallized and determined a 1.9 Å resolution crystal structure of *E. coli* Ubl_3_. The structure revealed the expected array of three β-grasp domains, with the first two domains tightly associated with one another through a predominantly hydrophobic interface burying ∼500 Å^2^ of surface area per domain. The interface between the second and third β-grasp domains is much smaller (280 Å^2^ per domain) and polar, implying flexibility (**Figure 2a-b**). All three domains adopt the β-grasp fold, with five antiparallel β-strands embracing a single short a-helix (**Figure 2c-d**). In β-grasp domains 2 and 3, the short a-helix is buttressed by an additional two-stranded antiparallel β-sheet comprising one strand from the loop linking β-strands S4 and S5, and a second strand just upstream of the a-helix (which is itself inserted between β-strands S2 and S3). Within the crystallographic asymmetric unit, we observed a dimeric assembly mediated by an antiparallel arrangement of the first two β-grasp domains, burying ∼1,100 Å of mostly polar surface area per protomer (**Figure 2b**). The dimer is apparently stabilized by two symmetrically-bound divalent cations (built as Ca^2+^ based on coordination geometry and refined difference electron density), which are each coordinated by three main-chain carbonyl groups and a glutamate side-chain from one protomer (E136), plus an aspartate side-chain from the second protomer (D73; **Figure 2e**). Sequence alignments show that these residues are highly conserved across Ubl_3_ proteins (**Figure S3a-b**).

**Figure 2.**
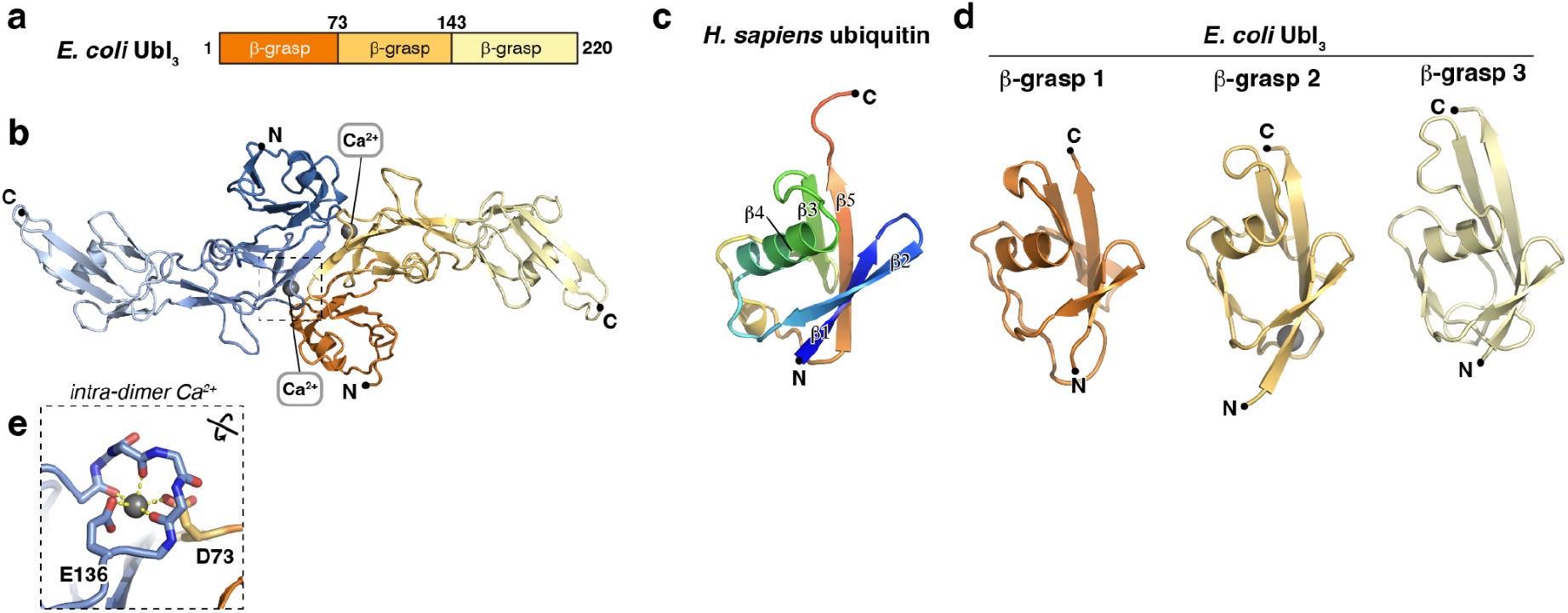
Crystal structure of *E. coli* Ubl_3_. **(a)** Domain structure of *E. coli* Ubl_3_. **(b)** Crystal structure of an *E. coli* Ubl_3_ dimer, with one protomer colored dark orange/light orange/light yellow and the second protomer colored dark blue/medium blue/light blue. Bound Ca^2+^ ions are shown as gray spheres. **(c)** Structure of *H. sapiens* ubiquitin (PDB ID 1UBQ) ^36^, colored as a rainbow from N-terminus (blue) to C-terminus (red) and with β-strands 1-5 labeled. **(d)** Structures of *E. coli* Ubl_3_ β-grasp domains 1-3, colored as in panel (b) and aligned with the structure of *H. sapiens* ubiquitin. **(e)** Close-up of the intra-dimer Ca^2+^ ion bound by *E. coli* Ubl_3_ indicated by a dotted box in panel (b). See **Figure S3a** for a sequence alignment of Ubl_3_ β-grasp domains 1-2 showing conservation of this site.

We next crystallized and determined a 3.0 Å resolution crystal structure of *M. brachiatum* Ubl_3_. In the structure, we observed only the first two of three predicted β-grasp domains; the third domain is likely to be flexible and disordered (**Figure 3a-b, Figure S4a-d**). Like *E. coli* Ubl_3_, *M. brachiatum* Ubl_3_ forms a symmetric homodimer through association of β-grasp domains 1 and 2, and the dimer symmetrically binds two ions (again built as Ca^2+^), which are coordinated identically to those bound by *E. coli* Ubl_3_, by E148 and D85 (**Figure 3b** *inset*).

**Figure 3.**
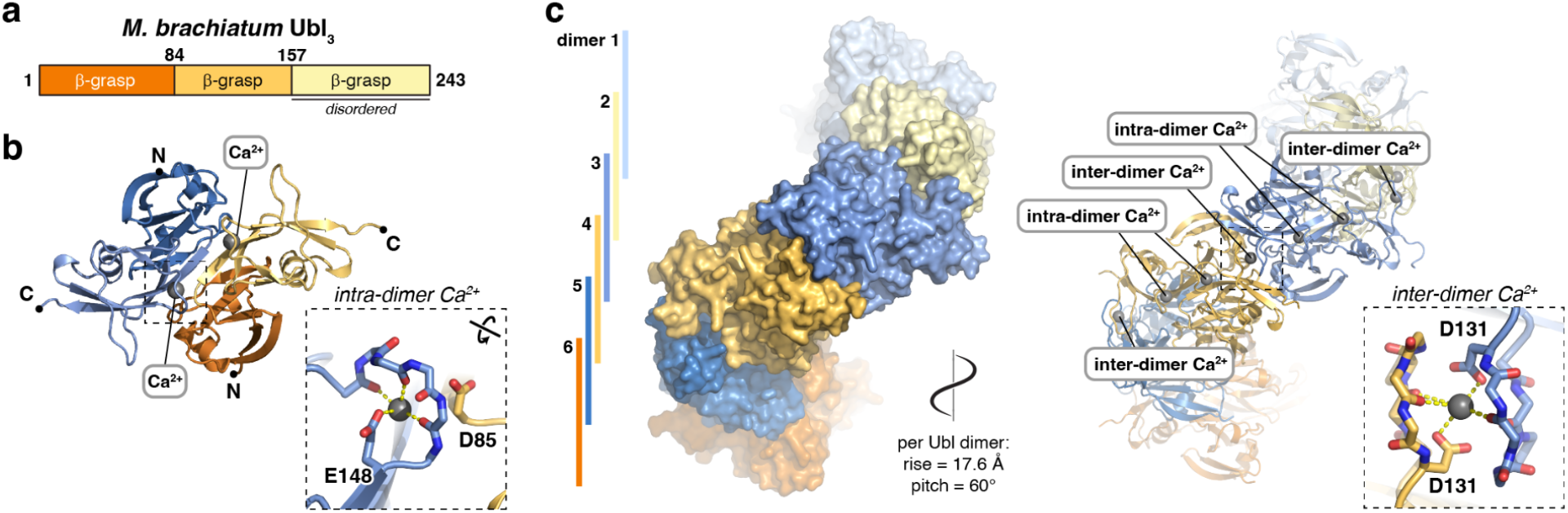
Crystal structure of *M. brachiatum* Ubl_3_. **(a)** Domain structure of *M. brachiatum* Ubl_3_. **(b)** Crystal structure of an *M. brachiatum* Ubl_3_ dimer, with one protomer colored dark orange/light orange and the second protomer colored dark blue/medium blue (β-grasp domain 3 is disordered and not shown). Bound Ca^2+^ ions are shown as gray spheres. *Inset:* Close-up of the intra-dimer Ca^2+^ ion. See **Figure S4** for detailed views of individual β-grasp domains in each bacterial Ubl_3_ protein. **(c)** Structure of a *M. brachiatum* Ubl_3_ filament assembled by a combination of crystallographic and non-crystallographic symmetry. Six Ubl_3_ dimers (three asymmetric units related by a P3_1_ screw axis) are depicted in alternating blue and orange. Bound Ca^2+^ ions are shown as spheres in the cartoon depiction at right and labeled according to whether they link protomers within a dimer (intra-dimer) or link protomers in different dimers (inter-dimer). *Inset:* Close-up of the inter-dimer Ca^2+^ ion.

Within the crystallographic asymmetric unit, two *M. brachiatum* Ubl_3_ dimers pack against one another and are related by a 60° right-handed rotation and an ∼18 Å translation along the crystallographic c axis. Further, these crystals adopt space group P3_1_21, which contains a screw axis along the crystallographic c axis with a right-handed twist of ∼120° and a rise of 35.2 Å (105.5 Å c axis divided by three) per asymmetric unit. Combining non-crystallographic and crystallographic symmetry gives an apparent helical filament of *M. brachiatum* Ubl_3_ dimers, with a right-handed helical twist of 60° and a rise of 17.6 Å (105.5 Å divided by six) per dimer (**Figure 3c**). We observe an additional ion symmetrically bound between each Ubl_3_ dimer pair, coordinated by two main-chain carbonyl groups and an aspartate side-chain (D131) from each of the two protomers involved (**Figure 3c** *inset*). Sequence alignments show that the acidic residues involved in both intra-dimer and inter-dimer Ca^2+^ ion coordination are highly conserved across Ubl_3_ proteins, implying that these sites are functionally relevant (**Figure S3a-b**).

### Ubl_3_ proteins oligomerize in the presence of Ca^2+^

Our structures of *E. coli* Ubl_3_ and *M. brachiatum* Ubl_3_ suggest that these proteins bind ions at conserved sites within and between Ubl_3_ dimers, and that ion binding could mediate assembly of dimers and helical filaments. When we purified *Citrobacter* Ubl_3_, we found that the protein eluted from a size exclusion column in two peaks: one at the void volume (suggestive of a large oligomer), and one at the position expected for a monomer (**Figure S5a**). We pooled and concentrated fractions for the oligomer peak and subjected the protein to inductively coupled plasma mass spectrometry (ICP-MS) to measure metal ion content. We measured ∼0.2 molar equivalents of calcium, ∼0.05 molar equivalents of magnesium, and vanishingly small amounts of manganese and iron (see **Materials and Methods**). These data support our assignment of ion densities in the crystal structures of *M. brachiatum* and *E. coli* Ubl_3_ as Ca^2+^.

To further explore the role of divalent cations in Ubl_3_ oligomerization, we performed size exclusion chromatography coupled to multi-angle light scattering (SEC-MALS) on three Ubl_3_ proteins. We incubated purified *E. coli* Ubl_3_, *M. brachiatum* Ubl_3_, and *Citrobacter* Ubl_3_ in a buffer containing EDTA to chelate any bound cations, then performed SEC-MALS (**Figure 4a-c**). In all three cases, we observed predominantly monomeric Ubl_3_, with *Citrobacter* Ubl_3_ migrating as two peaks: one representing a monomer and a second between the expected molecular weight of a monomer and a dimer (**Figure 4c**). We next removed EDTA by buffer-exchange and performed SEC-MALS in a buffer containing Mg^2+^ (**Figure 4d-f**). In this condition, *E. coli* Ubl_3_ and *M. brachiatum* Ubl_3_ migrated as monomers (**Figure 4d-e**), while *Citrobacter* Ubl_3_ shifted to a predominantly dimeric form (**Figure 4f**). Finally, we performed SEC-MALS in a buffer containing Ca^2+^ (**Figure 4g-i**). Both *M. brachiatum* Ubl_3_ and *Citrobacter* Ubl_3_ migrated in the void volume of the size exclusion column, forming large assemblies in the MDa size range (**Figure 4h-i**). *E. coli* Ubl_3_ precipitated from solution in the presence of 5 mM Ca^2+^, but we could perform SEC-MALS on the protein after incubation with 50 µM Ca^2+^. This analysis showed a mix of large oligomers in the MDa size range, plus apparent monomers and dimers (**Figure 4g**). Finally, we performed isothermal titration calorimetry (ITC) on *Citrobacter* Ubl_3_ to measure the affinity and stoichiometry of Ca^2+^ binding. This analysis showed that *Citrobacter* Ubl_3_ binds ∼4 Ca^2+^ ions per Ubl_3_ protomer, with the best fit to the data suggesting one high-affinity binding site (*K*_*d*_ ∼100 nM), and three lower-affinity binding sites (*K*_*d*_ ∼3 µM; **Figure S3c**).

**Figure 4.**
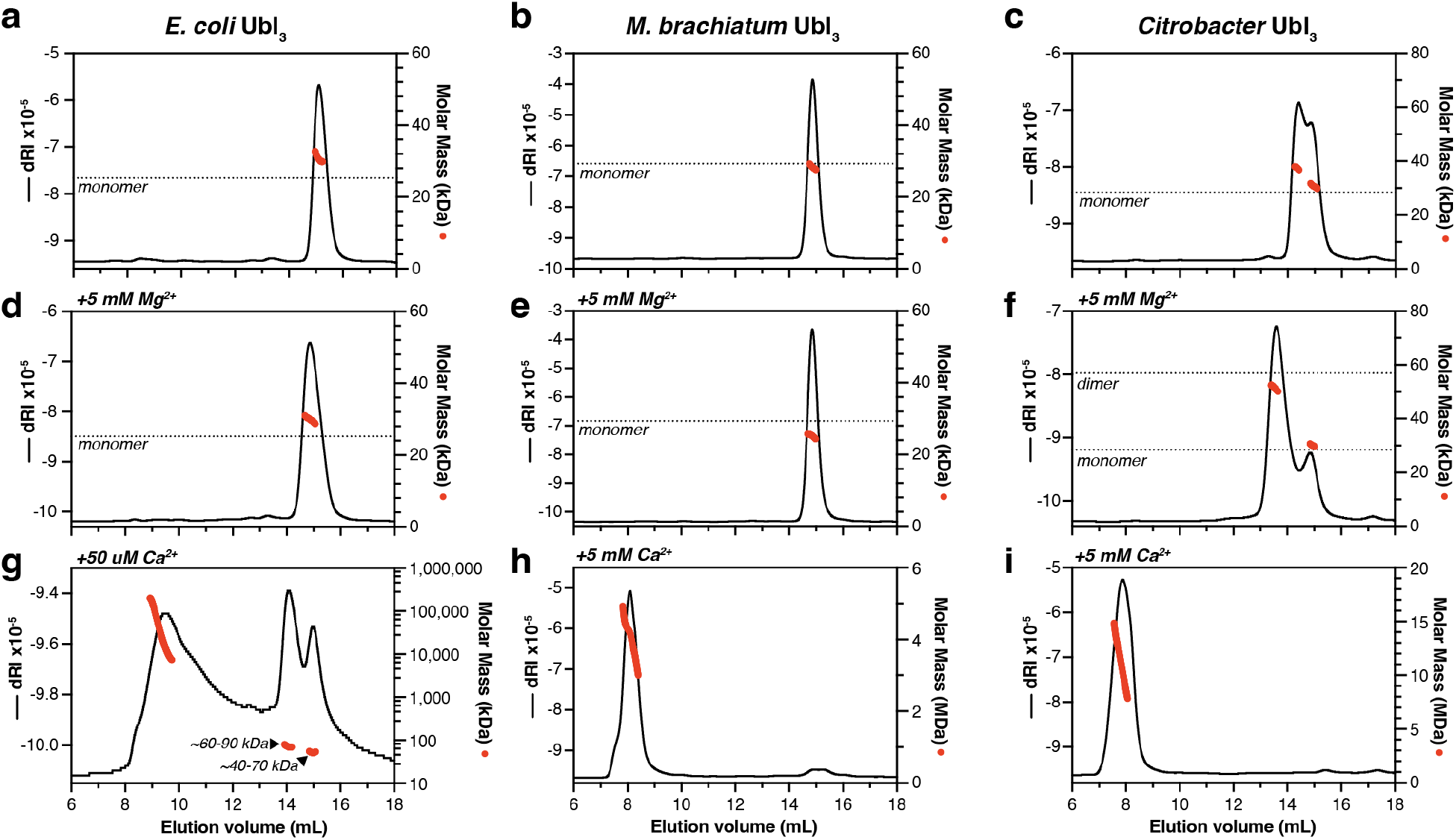
Ubl_3_ proteins oligomerize in the presence of Ca^2+^. **(a-c)** SEC-MALS analysis of *E. coli* Ubl_3_ (a), *M. brachiatum* Ubl_3_ (b), and *Citrobacter* Ubl_3_ (d) in the absence of divalent cations. Black lines indicate dRI (change in refractive index; left Y axis) and red dots indicate measured molar mass (right Y axis; shown in logarithmic scale). The expected molar mass of a monomer species is shown as a dotted line. **(d-f)** SEC-MALS analysis of *E. coli* Ubl_3_ (d), *M. brachiatum* Ubl_3_ (e), and *Citrobacter* Ubl_3_ (f) in the presence of 5 mM MgCl_2_. The expected molar mass of monomer and dimer species are shown as dotted lines. **(g-i)** SEC-MALS analysis of *E. coli* Ubl_3_ in the presence of 50 µM CaCl_2_ (g), *M. brachiatum* Ubl_3_ in the presence of 5 mM CaCl_2_ (h), and *Citrobacter* Ubl_3_ in the presence of 5 mM CaCl_2_ (i).

### Ubl_3_ proteins form helical filaments in the presence of Ca^2+^ ions

To determine the molecular basis for Ca^2+^-mediated Ubl_3_ oligomerization, we prepared cryoelectron microscopy (cryoEM) grids of oligomerized *Citrobacter* Ubl_3_ and *M. brachiatum* Ubl_3_. For *Citrobacter* Ubl_3_, we used a fraction eluting in the void volume of a size exclusion column upon initial purification of the protein from *E. coli* expression without EDTA addition (**Figure S5a**). For *M. brachiatum* Ubl_3_, we added 5 mM CaCl_2_ to purified monomeric Ubl_3_, incubated 60 minutes at 4°C, then prepared cryoEM grids. Initial images of both samples showed filaments ∼20 nm in diameter (**Figure 5a, Figure S6a**). We collected a full cryoEM dataset for each sample and determined the structures using helical reconstruction methods, resulting in a 2.4 Å resolution structure of *Citrobacter* Ubl_3_ (**Figure S5**) and a 3.1 Å resolution structure of *M. brachiatum* Ubl_3_ (**Figure S6**).

**Figure 5.**
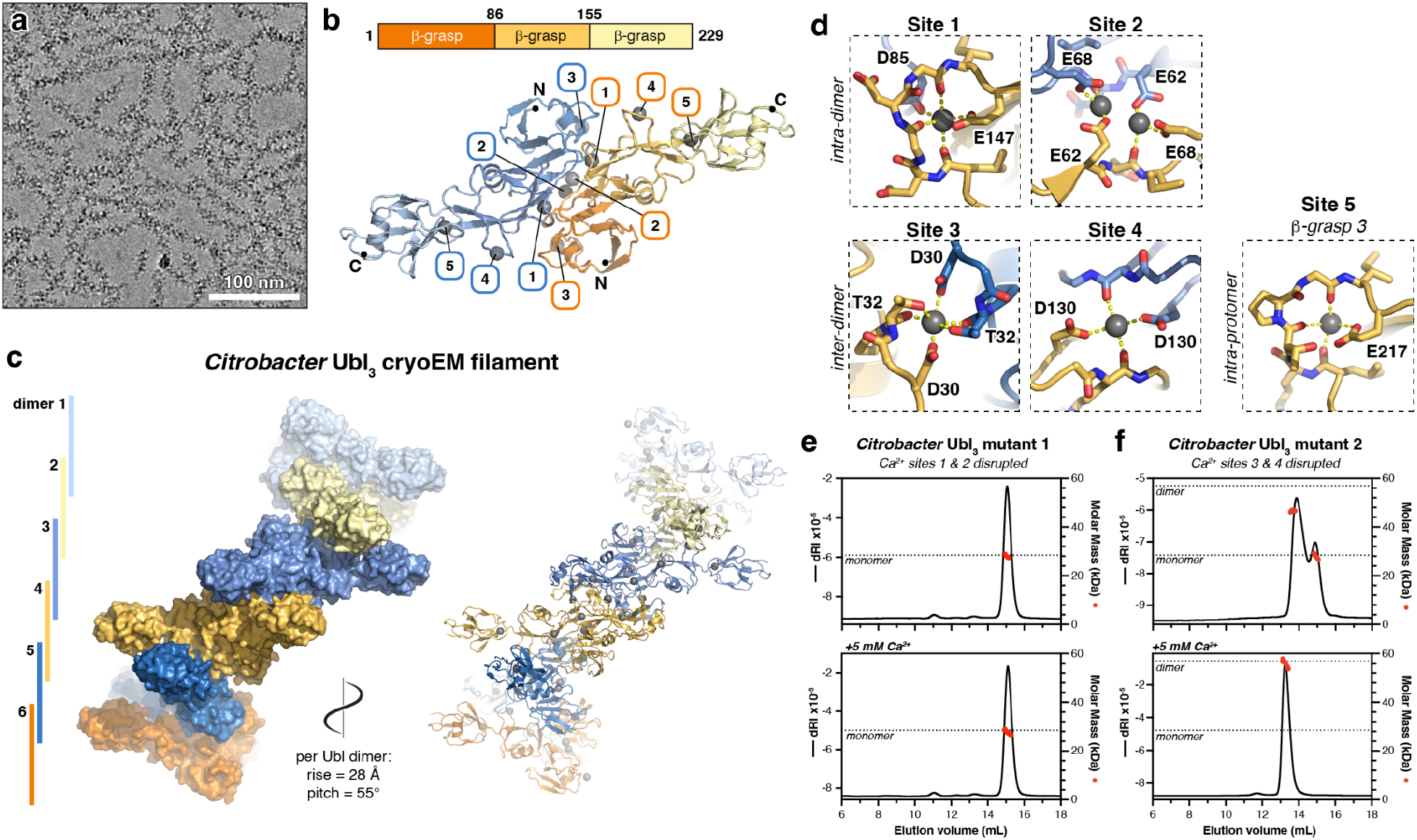
CryoEM structure of a *Citrobacter* Ubl_3_ filament. **(a)** CryoEM micrograph of *Citrobacter* Ubl_3_ filaments (see **Figure S5a** for size exclusion chromatography). **(b)** *Top:* Domain structure of *Citrobacter* Ubl_3_. *Bottom:* CryoEM structure of a *Citrobacter* Ubl_3_ dimer (see **Figure S5**) with one protomer colored dark orange/light orange/light yellow and the second protomer colored dark blue/medium blue/light blue. Bound Ca^2+^ ions are shown as gray spheres and labeled 1-5. **(c)** Closeup views of Ca^2+^ sites 1-5. Sites 1 and 2 link Ubl_3_ protomers within a dimer (intra-dimer), and sites 3 and 4 link Ubl_3_ protomers in different dimers (inter-dimer). Site 5 is coordinated entirely by residues in β-grasp domain 3. **(d)** CryoEM structure of the *Citrobacter* Ubl_3_ filament (see **Figure S5**), showing six Ubl_3_ dimers (alternating blue and orange). Bound Ca^2+^ ions are shown as spheres in the cartoon depiction at right. **(e)** SEC-MALS analysis of *Citrobacter* Ubl_3_ mutant 1 (E62A/E68A/D85A/E147A), designed to disrupt intra-dimer Ca^2+^ sites 1 and 2. Top panel shows analysis in the absence of divalent cations, and bottom panel shows analysis in the presence of 5 mM CaCl_2_. The expected molar mass of a monomer species is shown as a dotted line. **(f)** SEC-MALS analysis of *Citrobacter* Ubl_3_ mutant 2 (D30A/D130A), designed to disrupt inter-dimer Ca^2+^ sites 3 and 4. Top panel shows analysis in the absence of divalent cations, and bottom panel shows analysis in the presence of 5 mM CaCl_2_. The expected molar masses of monomer and dimer species are shown as a dotted line.

The cryoEM structure of *Citrobacter* Ubl_3_ reveals a helical filament with Ubl_3_ homodimers assembled identically to the dimers observed in our crystal structures of both *E. coli* and *M. brachiatum* Ubl_3_ proteins (**Figure 5b, S3e-g**), stacked into a helical filament with a right-handed helical twist of 55° and a rise of 28 Å per Ubl_3_ dimer (**Figure 5c**). Ubl_3_ dimer-dimer interactions are nearly identical to that observed in the crystal structure of *M. brachiatum* Ubl_3_, with β-grasp domains 1 and 2 highly ordered and mediating filament formation, and β-grasp domain 3 extending outward from the filament. Despite the flexibility of β-grasp domain 3, we could use 3D variability analysis and local refinement to generate a 2.7 Å resolution map showing all three β-grasp domains, enabling us build and refine a model of the full-length protein (**Figure 5b, Figure S3e-f, Figure S5h-i**).

The cryoEM structure of *M. brachiatum* Ubl_3_ reveals a helical filament with a right-handed helical twist of 63° and a rise of 30 Å per Ubl_3_ dimer, distinct from the parameters in our crystal structure of the same protein (right-handed twist of 60° and rise of 17.6 Å per Ubl_3_ dimer) (**Figure S6f**). The packing of Ubl_3_ dimers in both structures is equivalent, with small conformational changes at dimer-dimer interfaces resulting in a significantly more extended filament as imaged by cryoEM versus x-ray crystallography. As in our x-ray crystal structure, our cryoEM structure of *M. brachiatum* Ubl_3_ revealed no evidence of β-grasp domain 3, likely due to flexibility of the linker between β-grasp domains 2 and 3.

Close inspection of the cryoEM maps of both *Citrobacter* and *M. brachiatum* Ubl_3_ revealed the presence of multiple ordered Ca^2+^ ions bound to each Ubl_3_ protomer (**Figure 5d, Figure S3, Figure S4e**). We observed five Ca^2+^ ions bound to each *Citrobacter* Ubl_3_ protomer, three of which were also observed in the *M. brachiatum* Ubl_3_ cryoEM structure. Site #1 is equivalent to the intra-dimer Ca^2+^ ion we observed in our crystal structures of *E. coli* and *M. brachiatum* Ubl_3_. In *Citrobacter* Ubl_3_, this Ca^2+^ ion is coordinated by two main-chain carbonyl groups (residues 143 and 145) and the side-chain of a glutamate (E147) of one protomer, and the side-chain of an aspartate (D85) on the second protomer (**Figure 5d**). In the *M. brachiatum* Ubl_3_ cryoEM structure, site #1 involves three main-chain carbonyl groups and the side-chain of E148 from one protomer, and D85 from the second protomer is nearby (**Figure S6g**). A second intra-dimer Ca^2+^ ion (site #2) in *Citrobacter* Ubl_3_ is coordinated by the side chains of two glutamate residues on opposite protomers (E62 and E68). The residues at site #2 are less well conserved in Ubl_3_ proteins than the site #1 residues (**Figure S3**), and we did not observe this ion in either the *E. coli* or *M. brachiatum* Ubl_3_ crystal structures, or the *M. brachiatum* Ubl_3_ cryoEM structure.

We observe two inter-dimer Ca^2+^ ions in the *Citrobacter* Ubl_3_ helical filament (**Figure 5d**). The Ca^2+^ ion at site #3 is symmetrically coordinated by an aspartate side chain (D30) and both the main-chain carbonyl and side-chain hydroxyl of a threonine (T32) on Ubl_3_ protomers from two different dimers. Site #4 is also symmetric, and involves a main-chain carbonyl (residue 125) and an aspartate side chain (D130) side chain from two Ubl_3_ protomers from two dimers. The residues involved in both sites #3 and #4 are conserved in Ubl_3_ proteins (**Figure S3**), and we observe Ca^2+^ ions bound to both sites in our cryoEM structure of *M. brachiatum* Ubl_3_. (**Figure S6g**). Overall, our structural data closely match our ITC results that showed *Citrobacter* Ubl_3_ binding ∼4 molar equivalents of Ca^2+^ ions (**Figure S3c**): we identify five distinct Ca^2+^ binding sites in *Citrobacter* Ubl_3_, with the two inter-dimer sites symmetrically bridging two protomers each, thereby yielding a total of four molar equivalents of Ca^2+^ binding this protein.

To directly link Ca^2+^ binding to Ubl_3_ oligomerization, we designed two multi-site mutants of *Citrobacter* Ubl_3_: mutant 1 (E62A/E68A/D85A/E147A) was designed to disrupt Ca^2+^ binding to intra-dimer sites #1 and #2, and mutant 2 (D30A/D130A) was designed to disrupt inter-dimer Ca^2+^ binding to sites #3 and #4. By SEC-MALS, we found that mutant 1 is monomeric in solution in both the absence and presence of Ca^2+^ ions, suggesting that Ca^2+^ binding at the intra-dimer sites #1 and #2 is critical for higher-order assembly (**Figure 5e**). Like wild-type *Citrobacter* Ubl_3_, mutant 2 migrated as two peaks in the absence of Ca^2+^, one representing a monomer and a second between the expected molecular weight of a monomer and a dimer (**Figure 5f**). In the presence of Ca^2+^, mutant 2 formed dimers but not higher-order assemblies. Thus, mutant 2’s ability to bind Ca^2+^ ions at the intra-dimer sites #1 and #2 enables dimer assembly, but its inability to bind Ca^2+^ ions at the inter-dimer sites #3 and #4 disrupts filament assembly. Overall, these data strongly implicate Ca^2+^ binding in filament formation by *Citrobacter* Ubl_3_.

## Discussion

It is now understood that bacteria possess a range of biochemical pathways related to eukaryotic ubiquitination, which mediate protein conjugation in a variety of contexts including antiphage immunity^7,8,10–13,15^. To date, Type II CBASS operons were demonstrated to bear homology to a noncanonical eukaryotic ubiquitination pathway acting in autophagy^10,11^, and Type I/II Bil (bacterial ISG15-like) operons were shown to be related to canonical eukaryotic ubiquitination pathways ^12^. Finally, two groups of Bub operons, previously termed 6E or DUF6527 operons^7,8,10^, encode predicted E1, E2, peptidase, and DUF6527 proteins in addition to Ubls with diverse predicted architectures^14^.

While bacterial Bil and Bub pathways encode Ubls with high structural similarity to eukaryotic ubiquitin, Ubl-target conjugation in bacteria is unlikely to constitute a degradation signal as it does in eukaryotic cells. Therefore, the functional consequences of Ubl-target conjugation in bacteria remains a key unanswered question. For one Type I Bil system, specific Ubl conjugation to phage central tail fiber proteins was shown to disrupt assembly of virions and also impair infectivity of fully-assembled virions^13^. The specific targets and the biological consequences of Ubl-target conjugation in Type II Bil and in Type I/II Bub are thus far unknown. These pathways may similarly function in antiphage immunity, or they may constitute stress response pathways that provide a selective advantage to their host in specific stress or pathogenic contexts.

A widespread group of bacterial Ubl proteins found in both Type II Bil and Type I/II Bub pathways is the Ubl_3_ family, whose members possess three ubiquitin-like β-grasp domains. We show here that Ubl_3_ proteins form helical filaments upon binding Ca^2+^ ions, and that the Ca^2+^ binding sites and the property of filament formation are conserved across diverse Ubl_3_ proteins.

We envision two possible scenarios for these filaments’ function in the context of stress response. In the first scenario, unperturbed cells possess Ubl_3_ filaments that cannot be used as a substrate for the pathways’ E1 and E2 proteins for target conjugation. Upon stress that reduces intracellular Ca^2+^, Ubl_3_ filaments would disassemble, freeing Ubl_3_ to be conjugated to target proteins. In a second scenario, Ubl_3_ proteins are monomeric in unperturbed cells, possibly pre-conjugated to target proteins. Upon stress that increases intracellular Ca^2+^, Ubl_3_ filaments would form and thereby sequester the conjugated target proteins. This scenario is conceptually similar to the known tendency of some metabolic proteins to form filaments in stress conditions that reversibly alter their activity^21^; in at least one case, filamentation of bacterial glutamine synthetase was shown to be regulated by metal ion binding^22^. In either scenario, we hypothesize that filament formation by Ubl_3_ proteins represents a mechanism to respond to stress conditions that alter intracellular Ca^2+^ concentration.

Bacterial Ubls show high diversity, possessing one to three β-grasp domains and N-terminal domains with unknown functions, some of which are predicted to mediate oligomerization. Just as bacterial ubiquitination-related pathways show high diversity and likely act in many stress-response contexts, bacterial Ubls likely have many roles that depend on their diverse architectures. Notably, a large proportion of bacterial Bub operons are associated with known DNA damage-responsive transcriptional regulators, including the WYL domain protein CapW/BrxR^23–25^ and the two-protein CapH+CapP system^26^ (**Table S1**). Future studies will likely need to move beyond antiphage immunity to explore the full range of adaptive benefits provided by these diverse, and as-yet mysterious, pathways.

## Materials and Methods

### Bioinformatics

To analyze bacterial Ubl diversity, Type II Bil operons identified by Chambers et al. ^12^ and DUF6527-containing operons identified by Millman et al. ^15^ were visually inspected in the IMG (Integrated Microbial Genomes & Microbiomes) database (https://img.jgi.doe.gov) for operon architecture and neighboring transcription factors. Structure predictions of Ubl proteins were performed by AlphaFold 2 ^17^, AlphaFold 3 ^18^, or ESMFold ^16^. Disorder/LLPS predictions were performed using catGRANULE (http://service.tartaglialab.com/new_submission/catGRANULE) ^27^, and coiled-coil predictions were performed using PairCoil2 (https://cb.csail.mit.edu/cb/paircoil2/) ^28^.

### Protein expression, purification, and characterization

All proteins used in this study are listed in **Table S2**. Codon-optimized genes encoding Ubl proteins without their C-terminally fused E2 domains (when present) into *E. coli* expression vectors encoding an N-terminal TEV protease-cleavable His_6_-tag (UC Berkeley Macrolab vector 2B-T, Addgene ID 29666) or an N-terminal TEV protease-cleavable His_6_-maltose binding protein tag (UC Berkeley Macrolab vector 2C-T, Addgene ID 29706). Vectors were transformed into *E. coli* Rosetta 2 pLysS (EMD Millipore), and 1 L cultures were grown at 37°C in 2XYT media to an OD_600_ of 0.6 before induction with 0.25 mM IPTG at 20°C for 16-18 hours. Cells were harvested by centrifugation, resuspended in buffer A (25 mM Tris-HCl pH 8.5, 300 mM NaCl, 5 mM MgCl_2_, 10% glycerol and 5 mM mercaptoethanol) containing 5 mM imidazole, then lysed by sonication (Branson Sonifier). Lysates were clarified by centrifugation, then supernatants were passed over a Ni-NTA Superflow column (Qiagen) in buffer A. The column was washed in wash buffer (buffer A containing 20 mM imidazole), then eluted in elution buffer (buffer A containing 400 mM imidazole). Eluates were concentrated by ultrafiltration (Amicon Ultra; EMD Millipore). For crystallography, His_6_-tagged proteins were cleaved with TEV protease (expressed and purified in-house from plasmid pRK793 (Addgene #8827)) ^29^ then passed over a HisTrap HP column (Cytiva) to remove His_6_-tagged TEV protease and uncleaved proteins. All proteins were finally passed over a Superdex 200 Increase size exclusion column (Cytiva) in size exclusion buffer (25 mM Tris-HCl pH 8.5, 300 mM NaCl, 5 mM MgCl_2_, 10% glycerol and 1mM DTT). Peak fractions were concentrated by ultrafiltration and stored at 4°C. For purification with EDTA treatment, eluates from the first Ni-NTA column were concentrated and 5 mM EDTA was added. After 16 hours at 4°C, the EDTA was removed by buffer exchange in centrifugal concentrators and the sample was passed over the size exclusion column in size exclusion buffer.

For characterization of oligomeric state by size exclusion chromatography coupled to multi-angle light scattering (SEC-MALS), 100 µl of purified proteins at a concentration of 5 mg/ml were injected onto a size exclusion column (Superdex 200 Increase 10/300 GL, Cytiva) in size exclusion buffer, then light scattering and differential refractive index (dRI) profiles were collected using miniDAWN TREOS and Optilab T-rEX detectors (Wyatt Technology). SEC-MALS data were analyzed using ASTRA software version 8 and visualized with Prism version 10 (GraphPad Software). For analysis of divalent cations’ role in oligomerization, proteins were pre-incubated in size exclusion buffer plus supplemented with 5 mM EDTA, 5 mM MgCl_2_, or 5 mM CaCl_2_ for 30 minutes at 4°C, then passed over the size exclusion column in the same buffer.

### Crystallography

For crystallization of *E. coli* ZDHYS365 Ubl_3_, purified protein was subjected to surface lysine methylation by treating with borane (50 mM final concentration) and formaldehyde (100 mM final concentration) for 1 hour at 4°C, followed by addition of glycine (25 mM final concentration) to quench the reaction for 30 minutes, followed by buffer exchange by centrifugal concentration. Methylated protein at 10 mg/mL in crystallization buffer (25 mM Tris-HCl pH 8.5, 200 mM NaCl, 5 mM MgCl_2_, and 1 mM TCEP (tris(2-carboxyethyl)phosphine)) was mixed 1:1 with well solution containing 1 M LiCl, 0.1 M sodium acetate, and 30% PEG 6000 in sitting drop format. Crystals were harvested directly from the crystallization drop and frozen in liquid nitrogen. Diffraction data were collected at the Advanced Light Source (Lawrence Berkeley National Lab) beamline 5.0.1 on August 30, 2022 (collection temperature 100 K; x-ray wavelength 0.97741) (**Table S3**). Data were processed with XDS ^30^ and converted to structure factors with TRUNCATE ^31^. The structure was determined by molecular replacement in PHASER ^32^ using a predicted structure from AlphaFold 2 ^17^ as a search model. The model containing two protomers was manually rebuilt in COOT ^33^ and refined in phenix.refine ^34^ using positional and individual isotropic B-factor refinement. The final model has good geometry, with 98.77% of residues in favored Ramachandran space, 1.23% allowed, and 0% outliers. The overall MolProbity score is 1.07, and the MolProbity clash score is 2.12.

For crystallization of *M. brachiatum* Ubl_3_, purified protein was subjected to surface lysine methylation as above. Methylated protein at 12 mg/mL in the crystallization buffer was mixed 1:1 with a well solution containing 100 mM CAPS pH 10.5, 0.2 M Li_2_SO_4_, 1.2 M NaH_2_PO_4_, and 0.8 M K_2_HPO_4_ in sitting drop format. Crystals were harvested into a cryoprotectant solution containing 30% glycerol and frozen in liquid nitrogen. Diffraction data were collected at the Advanced Photon Source (Argonne National Lab) NE-CAT beamline 24ID-C on March 2, 2023 (collection temperature 100 K; x-ray wavelength 0.97911 Å) (**Table S3**). Data were processed with the RAPD data-processing pipeline, which uses XDS for data indexing and reduction, POINTLESS ^31^ for space group assignment, and AIMLESS ^31^ for scaling. The structure was determined by molecular replacement in PHASER using a predicted structure from AlphaFold 2 as a search model. Despite crystallizing full-length protein, only β-grasp domains 1 and 2 were visible in the model. The model containing four protomers was manually rebuilt in COOT and refined in phenix.refine using positional and individual isotropic B-factor refinement. The final model has good geometry, with 99.49% of residues in favored Ramachandran space, 0.51% allowed, and 0% outliers. The overall MolProbity score is 1.11, and the MolProbity clash score is 3.22.

### ICP-MS

For ICP-MS, we purified *Citrobacter* Ubl_3_ in size exclusion buffer lacking divalent cations, and concentrated fractions eluting from a size exclusion column at the void volume for analysis, to a final concentration of 25 mg/mL (0.956 mM). We diluted the protein 100-fold into 1% nitric acid in ultrapure water, then analyzed metal content using an iCAP RQ ICP-MS instrument (Thermo Fisher Scientific). We measured manganese at 0.102 ppb (parts per billion), iron at 0.415 ppb, magnesium at 11.120 ppb, and calcium at 84.458 ppb. After accounting for the 100X dilution, these measurements correspond to 0.000185 mM manganese, 0.000727 mM iron, 0.0463 mM magnesium, and 0.192 mM calcium.

### Isothermal Titration Calorimetry

For ITC, *Citrobacter* Ubl_3_ was purified with EDTA treatment, then passed over a size exclusion column in a buffer containing 25 mM Tris-HCl pH 8.5, 300 mM NaCl, and 1 mM NaN_3_, and fractions representing monomeric protein were pooled and concentrated. ITC was performed using an ITC-200 instrument (Malvern Panalytical), and four independent runs were performed at 25°C with 20-33 µM protein in the analysis cell and 300 µM CaCl_2_ in the syringe (40 × 3 µL injections with 120 seconds delay between each injection). Data were fit using a two-site binding model.

### Cryoelectron Microscopy of *Citrobacter* Ubl_3_

For structure determination of *Citrobacter* sp. RHBSTW-00271 Ubl_3_, fractions eluting in the void of a Superdex 200 Increase size exclusion chromatography columns (Cytiva) were pooled and concentrated to 1 mg/mL. Prior to use, Quantifoil copper 1.2/1.3 300 mesh grids were plasma cleaned for 12 seconds using a preset program in the Solarus II plasma cleaner (Gatan). A 3.5 µL protein sample was applied to the grid within the environmental chamber adjusted to 4°C temperature and approximately 95% humidity in a Vitrobot Mark IV (Thermo Fisher Scientific). After a 5-second incubation, grids were blotted with a blot force of 4 for 4 seconds, the sample was then plunge-frozen into liquid nitrogen-cooled liquid ethane.

Grids were mounteds into standard AutoGrids (Thermo Fisher Scientific) for imaging. An initial dataset was collected on a Talos Arctica transmission electron microscope (Thermo Fisher Scientific) operated at 200 kV and equipped with a Gatan K3 direct electron detector. Movies were collected at a magnification of 130,000x and a pixel size of 0.95 Å, with a total dose of 55 e-/Å^2^. A defocus range of -0.5 to -2.2 was used during the data collection. In total, 396 movies were used in the final data processing after applying CTF fitting and excessive motion criteria.

All data processing was performed using cryoSPARC version 4 ^35^. From the initial dataset, particle picking was performed using the blob picker with an estimated particle diameter of 200 Å. 189,755 initial particles were subjected to multiple rounds to 2D classification, yielding seven high-quality 2D classes from 53,925 particles that were used for template-based particle picking using the cryoSPARC Filament Tracer job. 207,260 particles were subjected to 2D classification and *ab initio* reconstruction, yielding an initial 3D volume from 101,204 particles. This volume was used to generate templates for a second round of particle picking using the Filament Tracer job. 189,327 particles were subjected to 2D classification, *ab initio* reconstruction, and helical refinement, yielding a final 3.4 Å resolution map.

The final dataset was collected on a Titan Krios G4 transmission electron microscope (Thermo Fisher Scientific) operated at 300 kV and configured for fringe-free illumination, equipped with a Falcon 4 direct electron detector with Selectris X energy filter. The microscope was operated in EFTEM mode with a slit width of 20 eV. Automated data acquisition was performed using EPU (Thermo Fisher Scientific). Movies were collected at a magnification of 130,000x and a pixel size of 0.935 Å, with a total dose of 45 e-/Å^2^. A defocus range of -0.5 to -2.2 was used during the data collection. In total, 1,851 movies were used in the final data processing after applying CTF fitting and excessive motion criteria (**Table S4**).

Particle picking in the final dataset was performed using templates generated from the 3.4 Å map from the initial dataset, in the Filament Tracer job. The filament diameter was set to 200 Å and the separation distance between picks was set to 0.14 diameters, enabling picking of adjacent asymmetric units along the helical filament. Particles were picked with a 320×320 pixel box size and Fourier cropped to 80×80 pixels for initial processing. 1,634,973 particles were subjected to four successive rounds of 2D classification (in rounds 1 and 2, the data were split into three batches), yielding a final set of 892,949 particles. *Ab initio* reconstruction was performed with three models, and the best model contained 640,938 particles. Particles were re-extracted at 320×320 pixels with re-centering (yielding 629,161 particles) and used for helical refinement with D1 symmetry (two-fold rotational symmetry oriented 90° from the helical axis). Helical symmetry was not applied during refinement as initial particle picking used a single-asymmetric-unit step. Final helical parameters are a right-handed helical twist of 55.78° per asymmetric unit, and a helical rise of 28.4 Å per asymmetric unit. The final refinement using the Helical Refinement job type resulted in a 2.6 Å resolution map revealing β-grasp domains 1 and 2, which participate in helical packing. To resolve the flexible third β-grasp domain, particles were symmetry-expanded (using D1 symmetry) then subjected to local refinement using a mask covering four protomers. The resulting 2.4 Å resolution map was subjected to 3D variability analysis and clustering, resulting in one class with strong density for the third β-grasp domain. This class (containing 148,697 particles) was refined to 2.7 Å resolution (**Table S4**).

For modeling, initial models of each β-grasp domain generated by AlphaFold 2 were manually placed, then the model was manually rebuilt in COOT with manual addition of ordered Ca^2+^ ions. The model was refined in phenix.refine using positional and individual isotropic B-factor refinement. For global refinement with four Ubl_3_ protomers, strict non-crystallographic symmetry restraints were applied during refinement for all protein atoms (**Table S4**).

### Cryoelectron Microscopy of *M. brachiatum* Ubl_3_

For structure determination of *M. brachiatum* Ubl_3_, pooled fractions from the monomeric peak of EDTA-treated protein at 0.5 mg/mL were mixed with 5 mM CaCl_2_, incubated 60 minutes at 4°C, then a 3.5 µL protein sample was applied to a plasma-cleaned Quantifoil grid within the environmental chamber adjusted to 4°C temperature and approximately 95% humidity in a Vitrobot Mark IV (Thermo Fisher Scientific). After a 5-second incubation, grids were blotted with a blot force of 4 for 4 seconds, the sample was then plunge-frozen into liquid nitrogen-cooled liquid ethane.

Grids were mounteds into standard AutoGrids (Thermo Fisher Scientific) for imaging, and a 1074 movie dataset was collected on a Titan Krios G3 transmission electron microscope (Thermo Fisher Scientific) operated at 300 kV and equipped with a Gatan K3 direct electron detector. Movies were collected at a magnification of 130,000x and a pixel size of 0.95 Å, with a total dose of 50 e-/Å^2^. A defocus range of -1.2 to -2 was used during the data collection. In total, 1039 movies were used in the final data processing after applying CTF fitting and excessive motion criteria (**Table S4**).

All data processing was performed using cryoSPARC version 4. From the initial dataset, particle picking was performed using the Filament Tracer job with an estimated filament diameter of 120 Å. 1,303,971 initial particles were extracted with a box size of 400 pixels and Fourier cropped to 100 pixels. These particles were subjected to multiple rounds to 2D classification, yielding a dataset of 216,012 particles that were used for *ab initio* 3D reconstruction. 117,271 particles segregating to the best class were re-extracted at full resolution and refined using the Helical Refinement job with D1 symmetry, yielding a 3.22 Å map. This map was used to generate templates for a second round of particle picking using the Filament Tracer type, yielding a set of 495,217 particles. These were subjected to multiple rounds of 2D classification and 3D heterogeneous refinement before a final set of 151,090 particles were refined using the Helical Refinement type with D1 symmetry, yielding a final 3.08 Å resolution map (**Table S4**).

An initial model from the x-ray crystal structure was manually placed, then the model was manually rebuilt in COOT with manual addition of ordered Ca^2+^ ions. The model was refined in phenix.refine using positional and individual isotropic B-factor refinement. For global refinement with four Ubl_3_ protomers, strict non-crystallographic symmetry restraints were applied during refinement for all protein atoms (**Table S4**).

## Acknowledgements

The authors acknowledge funding from the National Institutes of Health (R35 GM144121 to K.D.C.) and from the Howard Hughes Medical Institutes Emerging Pathogens Initiative (to K.D.C.). L.R.C. is supported by the UCSD Molecular Biophysics Training Grant (T32 GM139795). The authors acknowledge the facilities of the UC San Diego cryo-EM facility, part of the Goeddel Family Technology Sandbox, along with the scientific and technical assistance of facility staff member Dr. Inga Kuschnerus. The authors acknowledge technical assistance from beamline staff at the Advanced Light Source at Lawrence Berkeley National Lab, and the Advanced Photon Source at Argonne National Lab. The Berkeley Center for Structural Biology is supported in part by the Howard Hughes Medical Institute. The Advanced Light Source is a Department of Energy Office of Science User Facility under Contract No. DE-AC02-05CH11231. The Pilatus detector on 5.0.1. was funded under NIH grant S10OD021832. The ALS-ENABLE beamlines are supported in part by the National Institutes of Health, National Institute of General Medical Sciences, grant P30 GM124169. This work is based upon research conducted at the Northeastern Collaborative Access Team beamlines, which are funded by the National Institute of General Medical Sciences from the National Institutes of Health (P30 GM124165). This research used resources of the Advanced Photon Source, a U.S. Department of Energy (DOE) Office of Science User Facility operated for the DOE Office of Science by Argonne National Laboratory under Contract No. DE-AC02-06CH11357.

