## Supplemental Information for "Structural diversity and oligomerization of bacterial ubiquitin-like proteins"

### Table S1. Ubiquitin-like protein architecture in Bil and Bub operons

*See separate Excel workbook*

### Table S2. Proteins used in this study

| Species | Operon type | Ubl type | Accession | Sequence |
|---|---|---|---|---|
| *Methylobacterium brachiatum* DSM19569 | Type II Bil | $Ubl_3$ | IMG 2928979547 | MHEHEHVEDRVAVQVFDENLNAKDVHLTDPVPTGRQIIKAA<br>GKHPVDDYAVLAWMPDNALRPLHLDETFDLRQHGVERILVA<br>PSDTLYRFFIDGQDQEWPVRGITGVVLKTLAGVDPAAFEVFL<br>VIPGDDDIRVEDHELFDLARKGVEHFQTVKRKAPAEHGIALK<br>VVVNGTETELKVHAGTPLRQVRTEALQKSGNVGRPEDEWQ<br>LKDEHGNPYDLNQTVAGVGLHDGDLVWLSLNAGVAGV |
| *Citrobacter* sp. RHBSTW-00271 | Type I Bub | $Ubl_3$ | IMG 2938140956 (Ubl-E2 fusion; E2 fragment not included) | MQDIQSQHHHRRFIEVADETLSFRQVVMEDSTPNGSQISAA<br>SGFKPDQMPVVLMLLPNGSLEDIRPDEVVDLSSEVRRFIVVE<br>SDRTYFFTIDGARLEWPCRFITGYSIRQLGDIGDNKKLLLERE<br>DEADLEVQNDQIIDLDGDGIERFISRKATWKLNIQGKEFTFDT<br>PTVVIRDAVIRAGLNPNQAWHIFLKVEGQPKVEKNIDDVIDLR<br>TPGIEKLRLTPKDVNNGEEPRATRRDFSLRPEDEHYLDEMG<br>YCWETRLVGNARWLIIHDYELPDGYNHHQVNLALLITSGYPV<br>NMLDMFYVYPPLVRVNGVNIPATEATVAIDSVAYQRWSRHR<br>SWNPEIDSVISQLAMADGCLQKEVGQ |
| *E. coli* ZDHYS365 | Type II Bub | $Ubl_3$ | NCBI WP_0530693 00.1 | MKNKFIKINDKIVEIDDLTPTGAQILLVAGVKDIVEYVLFQKLK<br>NGLLEEIRPEEKTTLDKEGVETFLMFNSDRTYRFTLNGKMFD<br>WGAPSLTGATVKSLAECDFDSNDVWLEMKNEKDKLITDHEH<br>IDLTQPNVERLYTQETSINIIVNAKMRVVNRRVISYWDVVHLA<br>YEHAENKETSIYSVDYAKGPISNPEGSMVDGQYVQLTDGMI<br>FYVTQTDKS |
| *Bradyrhizobium* sp. Ghvi | Type I Bub | Coiled-coil-$Ubl_1$ | IMG 2653856993 | MTEHDEAGAPTKREKELKAFRERQMRELREFEQRQKQELE<br>EFERQELEELKEFEERQHPYEIKIDRTEFKVTEHFLTGAQLR<br>ALPNPPIGPERDLFEVVPGGSDEKIADTQKVKMRDGLRFFTA<br>PAQINPGLL |
| *Bradyrhizobium nanningense* CCBAU 53390 | Type II Bil | Coiled-coil-$Ubl_1$ | IMG 2874628352 | MSAVHDNAGFGPERLANMSESGNKSAEQFDREAEFLKKEIE<br>GVNEVIRSEEKIKHDLEKREHELEEEARREREKHDHDHDHG<br>HLVRITMVVNGQPVVIEAEEKEKLTEIRQKALQETQNLAQPA<br>ENWEIKDEAGAVLDPEKRVGEYHFGKEVTLFLSLKAGVAG |
| *Pseudanabaena* sp. PCC 7367 | Type I Bub | $Ubl_2$ | IMG 2504678157 (Ubl-E2 fusion; E2 fragment not included) | MISTKKIQVQIDEKTYFVEDPVITGLQLLEKAEKRPSDEYLIFY<br>LLPGGQLEEIRLDETVDLRQMGIERFITWRSDRSFRLVIDGR<br>RFEWGIPIITGIQLKKLAGVDPKAYGIWLEVRGGEDRPIENAE<br>EVNLDAEGVERFFTGKKTTTEGQNAILLPSQDREYLKSRSLD<br>FEEIVDGSKCGVVFPGFPLPTQFDANQTDLLLILPSGYPDAP<br>PDMFFLDPWIKLRQGNCFPKAANQPYAFDGRSWQRWSRH<br>NREWRPGVDGIWTMLKRVEHALEVAA |
| *Bradyrhizobium* sp. WSM1417 | Type II Bub | $Ubl_2$ | IMG 2507506256 | MTDAKHEVRIHIDQKPYHSPNPTTGADLYELGHVAEGKVLY<br>KEVEGDHEDKLVRIDSPSIHLTEDEHFHSGDAPEKHYTIIVNT<br>DPVVVDHDVLTFEELVKIAYPVPPTGLDPEFTVSFEHAKSVP<br>HHGDLPPNGKVTVKKHGTIFDVDHTNRS |
| *Photobacterium chitinilyticum* BEI 247 | Type II Bub | DUF1508-$Ubl_1$ | IMG 2885240117 | MKGKFEIFQSSKNNEYYFRLKAAGNWEIILDSEGYTTKSSCL<br>NGIESVKENAQLEERFERLVAKNGEHYFNLKAGNGQVIGTS<br>EMYTRRQGMENGIHSVMKNAPDADIKDLTIDEPEHDKEFNII<br>VNGRPKTVTSKILTFEDIVKLAFSTIADGNSTIYTMTYKKNGG<br>NKPEGTLVTGDQIKIKSGVIFNVTATDKS |
| *Marinifilum flexuosum* isolate 1468 | Type II Bub | DUF1508-$Ubl_1$ | NCBI WP_2820164 92.1 | MNSYFTIKVGKDDQYYFNLKAGNHEIILQSEGYENKSGTLNG<br>IESVRLNSQIRSNFEIRYSKRNEPYFVL<br>KAPNGKIIGCSEMYSSVHAMENGIHSVMKNGNTPKIHDLTDD<br>DHEGKERQIVVNGRVKTWNQKYIEFKEL<br>VELAFGSYNDNPNTCYTITYTRGCSNKPQGSIVKGEEVKVK<br>PKMIFNVTATDKS |

**Table S3. Crystallographic data collection and refinement**

| | *E. coli* $Ubl_3$ | *M. brachiatum* $Ubl_3$ |
|---|---|---|
| **Data collection** | | |
| Data collection date | August 30, 2022 | March 2, 2023 |
| Beamline | ALS 5.0.1 | APS 24ID-C |
| Wavelength (Å) | 0.97741 | 0.97911 |
| Space group | $P3_221$ | $P3_121$ |
| Cell dimensions | | |
| *a*, *b*, *c* (Å) | 117.54, 117.54, 104.44 | 136.02, 136.02, 105.51 |
| α, β, γ (°) | 90, 90, 120 | 90, 90, 120 |
| Resolution (Å)* | 46.5-1.87 (1.91-1.87) | 117.8-3.04 (3.22-3.04) |
| $R_{merge}$ | 0.093 (2.232) | 0.204 (3.416) |
| $I / \sigma I$ | 16.1 (0.8) | 14.0 (0.9) |
| Completeness (%) | 100 (100) | 99.9 (99.9) |
| Redundancy | 9.5 (6.1) | 10.0 (10.4) |
| **Refinement** | | |
| Resolution (Å) | 46.46-1.87 | 117.80-3.04 |
| No. reflections | 69064 | 22023 |
| $R_{work}$ (%) | 18.16 (33.16) | 23.86 (39.41) |
| $R_{free}$ (%) | 21.27 (35.77) | 27.62 (39.99) |
| No. atoms | | |
| Protein (non-H) | 3526 | 4779 |
| Ligand/ion (non-H) | 28 | 31 ($Ca^{2+}$, $PO_4^-$) |
| Water | 537 | 53 |
| Hydrogen | 3511 | 0 |
| *B*-factors | | |
| Protein | 41.7 | 114.5 |
| Ligand/ion | 53.7 | 139.9 |
| Water | 45.5 | 98.5 |
| R.m.s. deviations | | |
| Bond lengths (Å) | 0.017 | 0.003 |
| Bond angles (°) | 1.399 | 0.57 |
| Validation | | |
| MolProbity score | 1.07 | 1.11 |
| Clashscore | 2.12 | 3.22 |
| Poor rotamers (%) | 1.30 | 0.40 |
| Ramachandran plot | | |
| Favored (%) | 98.77 | 99.49 |
| Allowed (%) | 1.23 | 0.51 |
| Disallowed (%) | 0 | 0 |
| PDB ID | 9CD2 | 8U38 |
| SBGrid Data Bank ID | 1115 | 1043 |

*Values in parentheses are for highest-resolution shell.

**Table S4. CryoEM data collection and refinement**

| | *Citrobacter* $Ubl_3$ (global) | *Citrobacter* $Ubl_3$ (local) | *M. brachiatum* $Ubl_3$ |
|---|---|---|---|
| **Data collection and processing** | | | |
| Magnification | 130,000 | | 105,000 |
| Voltage (kV) | 300 | | 300 |
| Electron exposure (e–/Å$^2$) | 45 | | 50 |
| Defocus range (μm) | -2.0 to -1.2 | | -2.0 to -1.2 |
| Pixel size (Å) | 0.935 | | 0.822 |
| Symmetry imposed | Helical and D1 | | Helical and D1 |
| Initial particle images | 1,634,973 | | 495,217 |
| Final particle images | 1,258,222 (D1 symmetry expanded) | 148,697 | 151,090 |
| Map resolution (Å; 0.143 FSC) | 2.43 | 2.73 | 3.08 |
| **Refinement** | *Global (4 protomers)* | *Local (1 protomer)* | *Global (4 protomers)* |
| Model resolution (Å; 0.143 FSC) | 2.4 | 2.7 | 3.1 |
| Model composition | | | |
| Non-hydrogen atoms | 4698 | 1704 | 4650 |
| Protein residues | 584 | 212 | 584 |
| Ligands ($Ca^{2+}$) | 14 | 5 | 10 |
| *B* factors (Å$^2$) | | | |
| Protein | 85.1 | 124.9 | 125.7 |
| Ligand | 108.1 | 110.0 | 90.2 |
| R.m.s. deviations | | | |
| Bond lengths (Å) | 0.002 | 0.004 | 0.002 |
| Bond angles (°) | 0.384 | 0.431 | 0.471 |
| Validation | | | |
| MolProbity score | 1.15 | 1.32 | 1.72 |
| Clashscore | 3.56 | 5.59 | 6.09 |
| Poor rotamers (%) | 0.76 | 1.05 | 3.23 |
| Ramachandran plot | | | |
| Favored (%) | 98.61 | 99.05 | 99.31 |
| Allowed (%) | 1.39 | 0.95 | 0.69 |
| Disallowed (%) | 0 | 0 | 0 |
| PDB ID | 9D59 | 9D5A | 9D5B |
| EMDB ID | EMD-46576 | EMD-46577 | EMD-46578 |

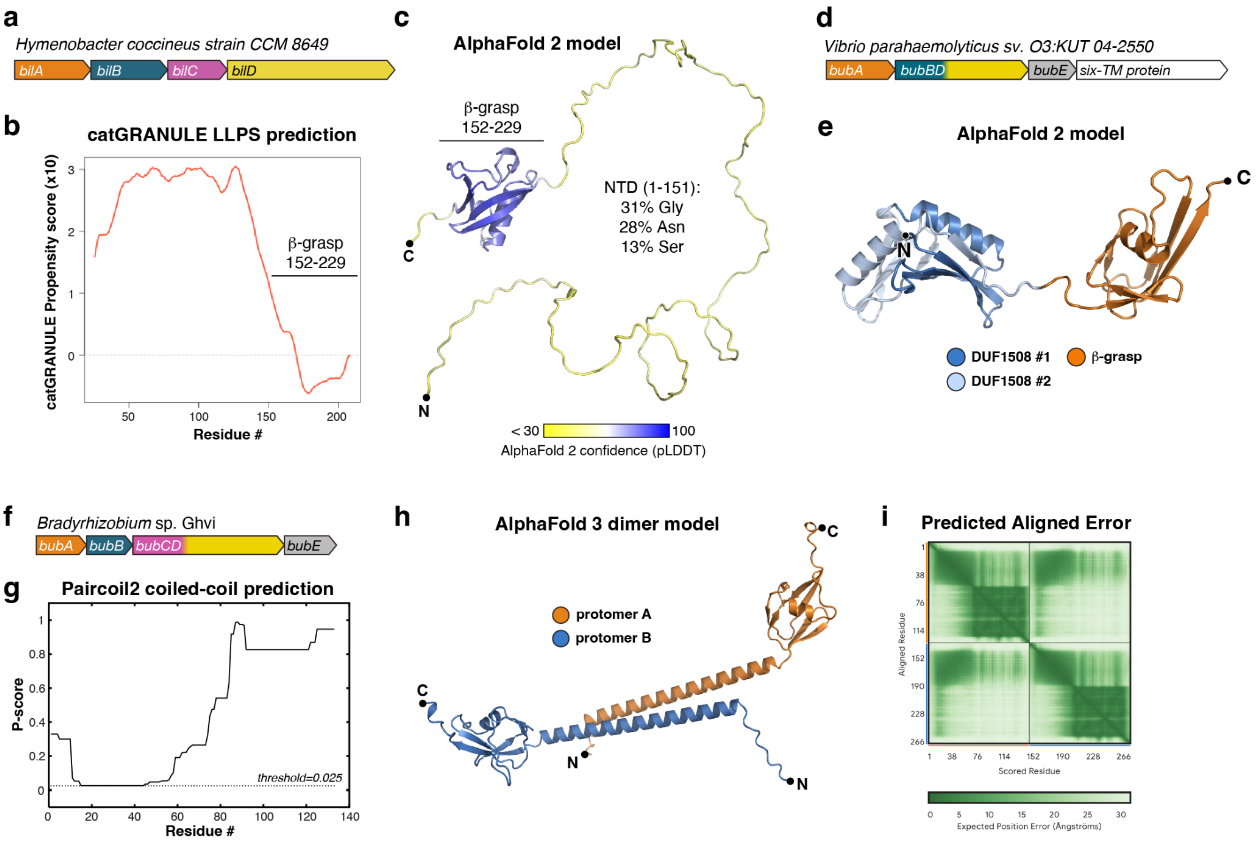

**Figure S1. Bacterial ubiquitin-like proteins show diverse architectures. (a)** Operon schematic of a *Hymenobacter* Type II Bil operon whose BilA protein (IMG Gene ID 2625034186) N-terminal domain is predicted to be disordered. **(b)** catGRANULE analysis of *Hymenobacter* BilA showing that its disordered N-terminal domain has a high propensity to undergo liquid-liquid phase separation. **(c)** AlphaFold 2 model of *Hymenobacter* BilA colored by confidence (pLDDT). The sequence composition of the disordered N-terminal domain is noted. **(d)** Operon schematic of a *Vibrio* Type II Bub operon whose BubA protein (IMG ID 2704903835) N-terminal domain is predicted to encode tandem DUF1508 domains. **(e)** AlphaFold2 model of *Vibrio* BubA, with DUF1508 domains colored dark/light blue and β-grasp domain dark orange. See **Figure S2c-d** for SEC-MALS analysis of two DUF1508-Ubl1 proteins. **(f)** Operon schematic of a *Bradyrhizobium* Type I Bub operon whose BubA protein (IMG ID 2653856993) N-terminal domain is predicted to form a coiled-coil. **(g)** PairCoil2 analysis of *Bradyrhizobium* BubA, showing a predicted coiled-coil domain at the N-terminus. **(h)** AlphaFold 3 model of a *Bradyrhizobium* BubA dimer, with one protomer colored dark orange and the second protomer colored dark blue. See **Figure S2e** for SEC-MALS analysis of this protein, and **Figure S2f** for SEC-MALS analysis of a second predicted CC-$Ubl_1$ protein. **(i)** AlphaFold 3 predicted aligned error plot for the *Bradyrhizobium* BubA dimer shown in panel (H), showing high confidence for interactions between dimer-related coiled-coil N-terminal domains.

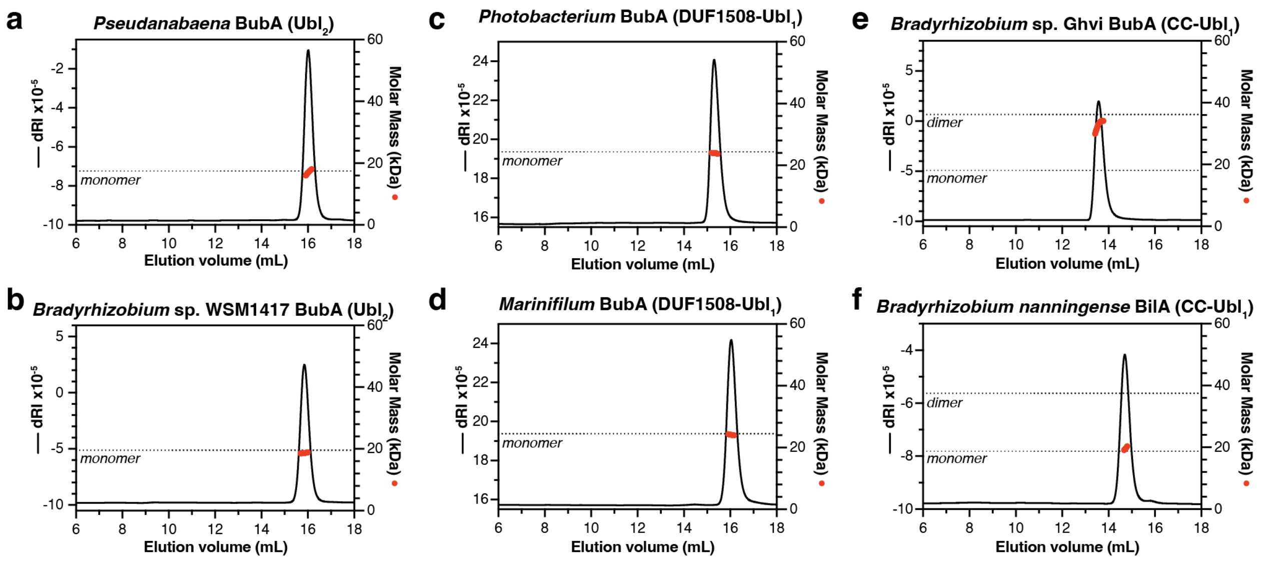

**Figure S2. SEC-MALS analysis of bacterial Ubl proteins. (a)** SEC-MALS analysis of *Pseudanabaena* sp. PCC 7367 BubA (see **Table S2** for accession numbers and protein sequences), which possesses two β-grasp domains. Differential refractive index (dRI; measuring protein concentration) is shown as a black line, and measured molar mass is shown as red circles. The predicted molar mass of a monomer is shown as a dotted line. **(b)** SEC-MALS analysis of *Bradyrhizobium* sp. WSM1417 BubA, which possesses two β-grasp domains. The predicted molar mass of a monomer is shown as a dotted line. **(c)** SEC-MALS analysis of *Photobacterium chitinilyticum* BEI 247 BubA, which possesses a single β-grasp domain and a predicted DUF1508 N-terminal domain. The predicted molar mass of a monomer is shown as a dotted line. **(d)** SEC-MALS analysis of *Marinifilum flexuosum* isolate 1468 BubA, which possesses a single β-grasp domain and a predicted DUF1508 N-terminal domain. The predicted molar mass of a monomer is shown as a dotted line. **(e)** SEC-MALS analysis of *Bradyrhizobium* sp. Ghvi BubA, which possesses a single β-grasp domain and a predicted coiled-coil N-terminal region. The predicted molar mass of a monomer and a dimer are shown as dotted lines. **(f)** SEC-MALS analysis of *Bradyrhizobium nanningense* CCBAU 53390 BilA, which possesses a single β-grasp domain and a predicted coiled-coil N-terminal region. The predicted molar mass of a monomer and a dimer are shown as dotted lines.

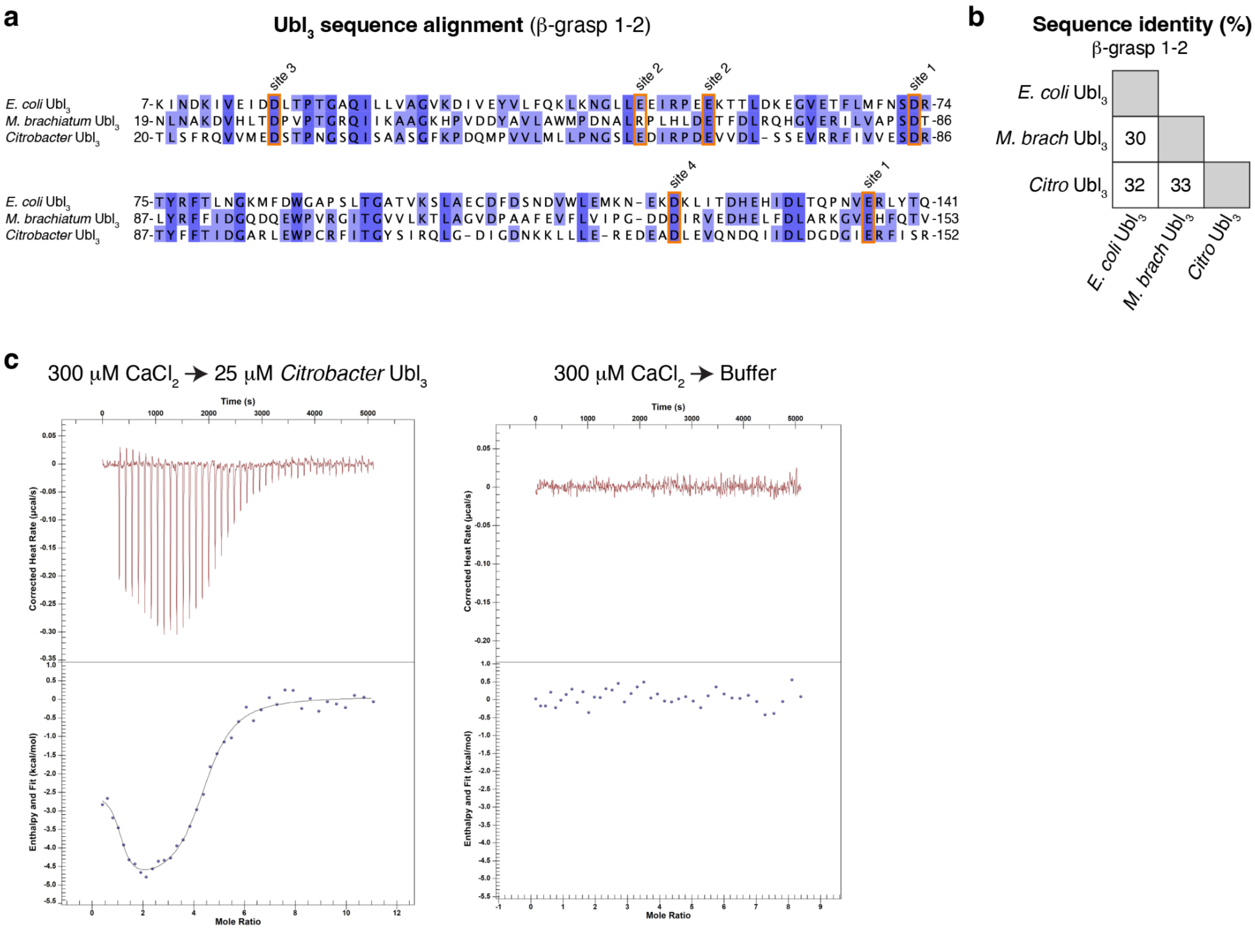

| | $K_d$ (μM) | N | ΔH (kcal/mol) | -TΔS (kcal/mol) |
|---|---|---|---|---|
| Site 1 | 0.09 +/- 0.05 | 1.2 +/- 0.2 | -2.1 +/- 0.5 | -7.7 +/- 0.2 |
| Site 2 | 2.8 +/- 0.4 | 3.4 +/- 0.1 | -4.6 +/- 0.5 | -3.0 +/- 0.4 |

**Figure S3. Sequence alignments and $Ca^{2+}$ binding of $Ubl_3$ proteins. (a)** Sequence alignment of β-grasp domains 1 and 2 from the three $Ubl_3$ proteins under study (see **Table S2**), with residues involved in $Ca^{2+}$ ion coordination outlined in orange. **(b)** Table of protein sequence identities for the three $Ubl_3$ proteins in panel (a). **(c)** Isothermal titration calorimetry for *Citrobacter* $Ubl_3$ binding $CaCl_2$. *Left:* Binding data (top) and fit curve (bottom); trace is representative of four independent trials. Right: No-protein control injection series. Bottom: Data from four independent trials was fit using a two-site binding model. N designates molar equivalent of ligand binding.

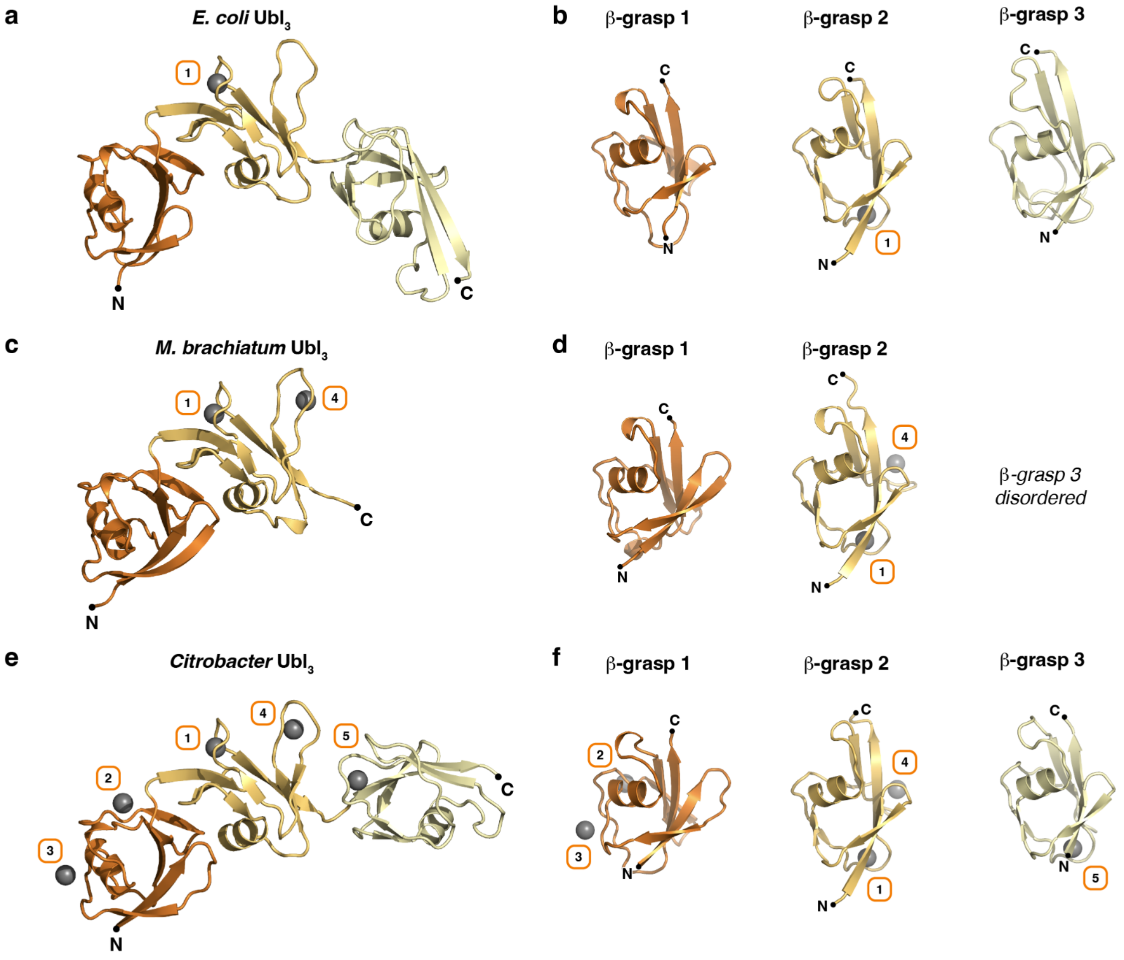

**g**

**Cα r.m.s.d. (Å)**
β-grasp 1-2

| | *E. coli* Ubl$_3$ | *M. brach* Ubl$_3$ | *Citro* Ubl$_3$ |
|---|---|---|---|
| *E. coli* Ubl$_3$ | | | |
| *M. brach* Ubl$_3$ | 0.81 | | |
| *Citro* Ubl$_3$ | 0.79 | 0.97 | |

**Figure S4. Structures of Ubl$_3$ proteins. (a)** Structure of an *E. coli* Ubl$_3$ monomer, with β-grasp domains 1-3 colored dark orange, light orange, and light yellow, respectively. Bound $Ca^{2+}$ ions are shown as gray spheres and labeled with the site number (according to *Citrobacter* Ubl$_3$ $Ca^{2+}$ site numbering). **(b)** Structures of *E. coli* Ubl$_3$ β-grasp domains 1-3. **(c)** Structure of an *M. brachiatum* Ubl$_3$ monomer. β-grasp domain 3 is disordered and not shown. **(d)** Structures of *M. brachiatum* Ubl$_3$ β-grasp domains 1-2. **(e)** Structure of a *Citrobacter* Ubl$_3$ monomer. **(f)** Structures of *Citrobacter* Ubl$_3$ β-grasp domains 1-3. **(g)** Table of structural similarity between three Ubl$_3$ proteins, spanning β-grasp domains 1 and 2. Cα r.m.s.d.: average root mean squared displacement between aligned alpha-carbon atoms. Comparison of *E. coli* and *Citrobacter* Ubl$_3$ β-grasp domain 3 shows higher divergence, with a 3.1 Å overall Cα r.m.s.d. in this domain.

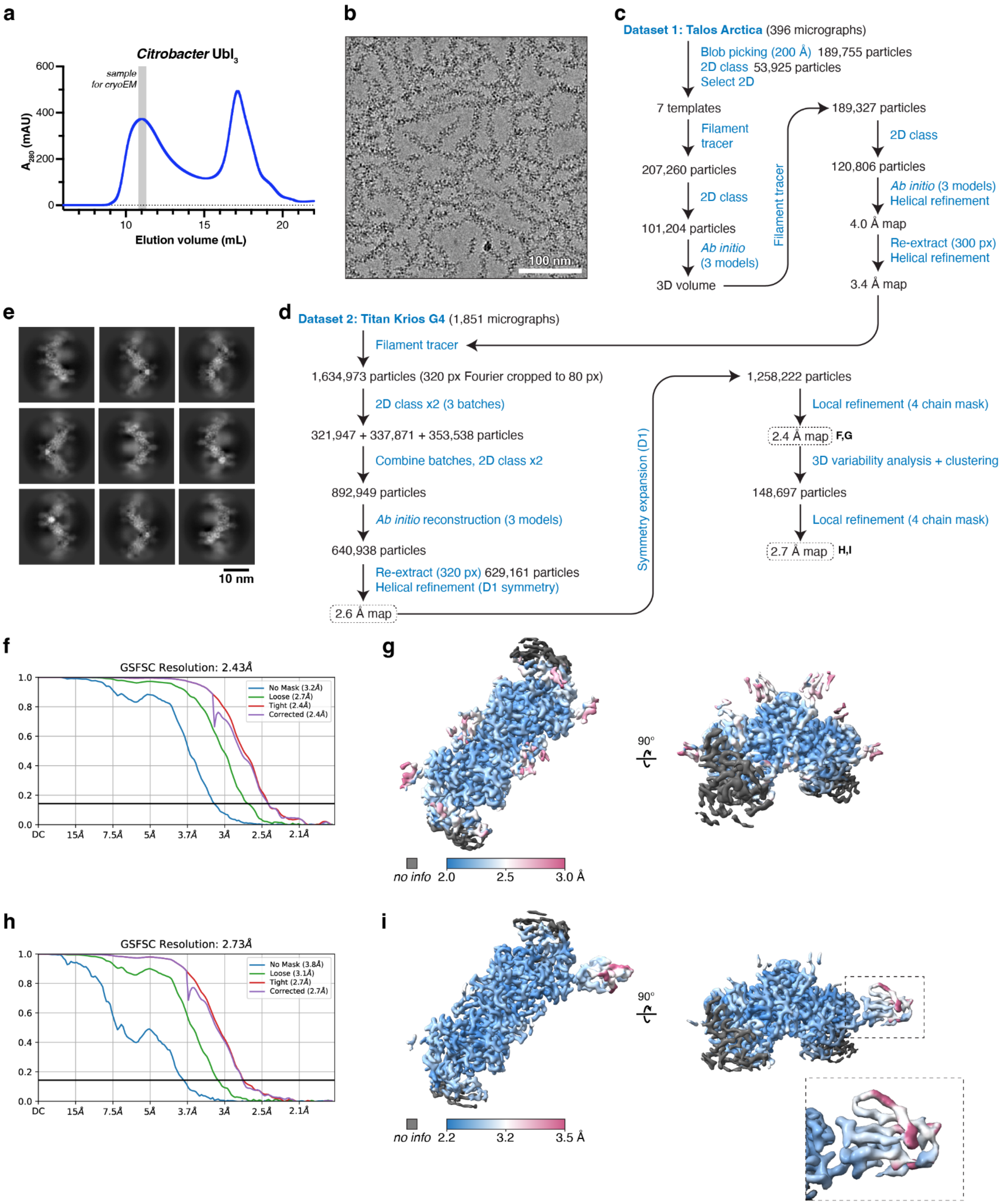

**Figure S5. CryoEM structure of a *Citrobacter* Ubl3 filament. (a)** Size exclusion chromatography elution profile of purified *Citrobacter* Ubl3, with the fraction used for cryoEM analysis highlighted in gray. **(b)** Raw cryoEM micrograph of *Citrobacter* Ubl3. **(c)** CryoEM structure determination workflow for preliminary dataset. **(d)** CryoEM structure determination workflow for final dataset. **(e)** Selected 2D classes from final dataset. **(f)** Fourier Shell Correlation graph for global refinement (four Ubl3 protomers masked). **(g)** Local resolution of final globally refined cryoEM map (four $Ubl_3$ protomers masked). **(h)** Fourier Shell Correlation graph for local refinement with particle subset selected from 3D variability analysis to reveal β-grasp domain 3 in one protomer (four $Ubl_3$ protomers masked). **(i)** Local resolution of local refinement with particle subset selected from 3D variability analysis to reveal β-grasp domain 3 in one protomer (four $Ubl_3$ protomers masked).

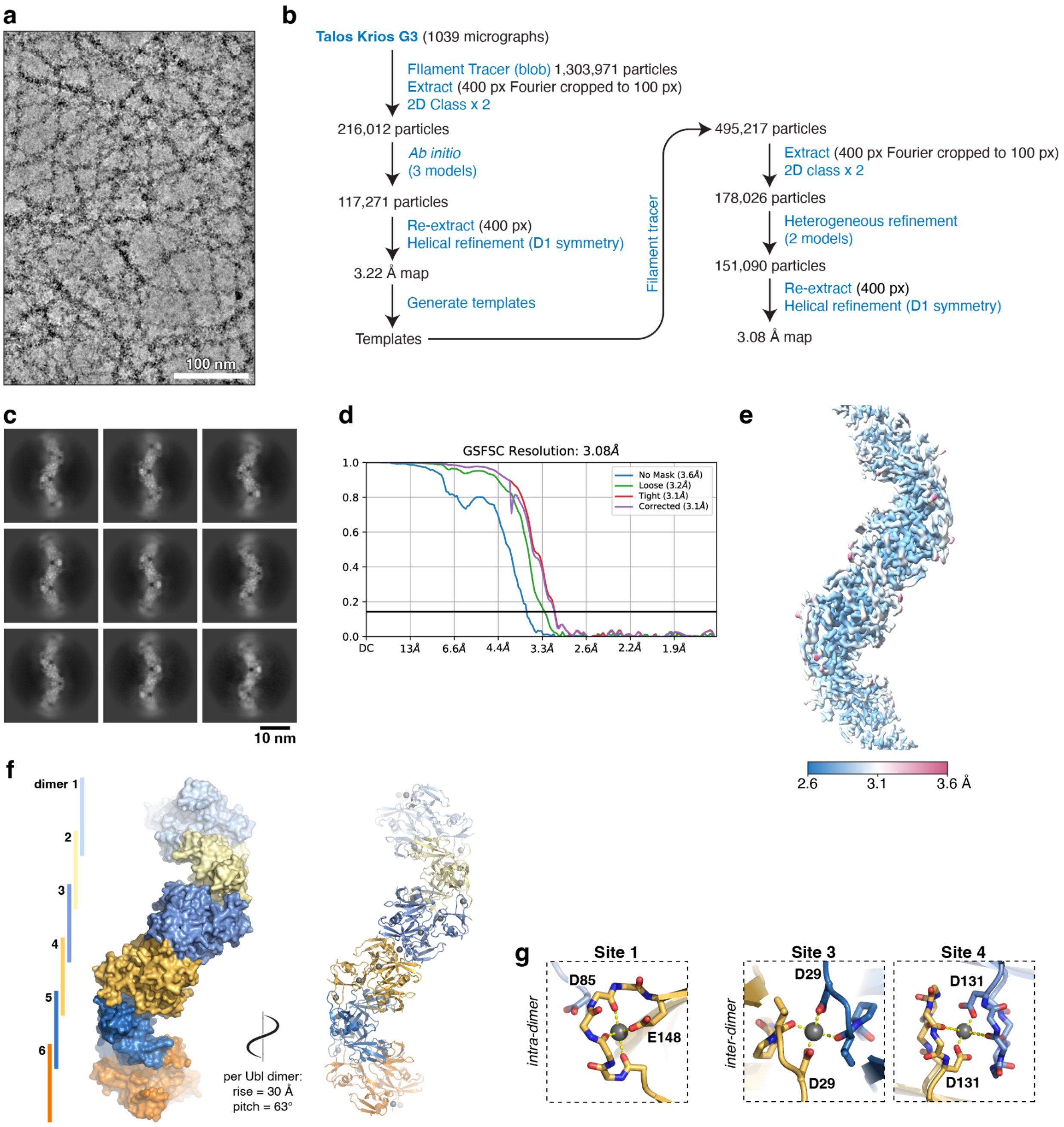

**Figure S6. CryoEM structure of an *M. brachiatum* Ubl$_3$ filament. (a)** Raw cryoEM micrograph of *M. brachiatum* Ubl$_3$ after incubation with 5 mM $CaCl_2$. **(b)** CryoEM structure determination workflow. **(c)** Selected 2D classes. **(d)** Fourier Shell Correlation graph for final refinement. **(e)** Local resolution of final refined cryoEM map. **(f)** Architecture of the *M. brachiatum* Ubl$_3$ filament, showing six Ubl$_3$ dimers in alternating blue and orange. Bound $Ca^{2+}$ ions are shown in gray on the cartoon diagram at right. **(g)** Closeup views of $Ca^{2+}$ ions bound at sites 1 (intra-dimer), 3 (inter-dimer), and 4 (inter-dimer).
